# *Toxoplasma gondii* PPM3C, a secreted protein phosphatase, affects parasitophorous vacuole effector export

**DOI:** 10.1101/2020.07.06.189308

**Authors:** Joshua A. Mayoral, Tadakimi Tomita, Vincent Tu, Jennifer T. Aguilan, Simone Sidoli, Louis M. Weiss

**Author notes:** Corresponding author: Louis M. Weiss, MD, MPH, Albert Einstein College of Medicine, 1300 Morris Park Avenue, Room 504 Forchheimer Building, Bronx, NY, 10461.

## Abstract

*Toxoplasma gondii* is a highly successful parasite that infects a significant portion of the human population. As an intracellular parasite, *T. gondii* thrives within many different cell types due to its residence in the parasitophorous vacuole, a specialized and heavily modified compartment in which parasites divide. Within this vacuole, numerous secreted proteins facilitate functions that optimize intracellular survival. We characterized one such protein, TgPPM3C, which is predicted to contain a domain belonging to the PP2C class of serine/threonine phosphatases and is secreted by both tachyzoites and differentiating bradyzoites into the vacuolar lumen. Genetic deletion of TgPPM3C established that parasites lacking this predicted phosphatase exhibit a minor growth defect *in vitro*, are avirulent during acute infection in mice, and form fewer cysts in mouse brain during chronic infection. A label-free phosphoproteomic approach was utilized to identify putative TgPPM3C substrates and demonstrated several secreted proteins with altered phosphorylation status in the absence of TgPPM3C. Altered phosphorylation status was seen in MYR1, a protein essential to the process of protein effector export from the parasitophorous vacuole into the host cell, and in GRA16 and GRA28, two exported effector proteins. Defects were seen in the export of GRA16 and GRA28, but not the effector TgIST, in the TgPPM3C knockout strain. Parasites lacking TgPPM3C also exhibited defects in host c-Myc induction, a process influenced by effector export. Phosphomimetic mutations of GRA16 serine residues recapitulated export defects, implicating de-phosphorylation as an important process in facilitating the export of GRA16. These findings provide an example of the emerging critical role that phosphatases play in regulating the complex environment of the *T. gondii* parasitophorous vacuole.

## INTRODUCTION

The intracellular parasite *Toxoplasma gondii* infects not only an impressive range of warm-blooded hosts, but can also infect any nucleated cell type within their host [1]. A large proportion of the human population, varying by region, is predicted to be infected based on seropositivity, posing a life-threatening risk in immunocompromised individuals [2]. During acute infection, the rapidly growing tachyzoite life stage disseminates robustly to various host tissues and cell types through repeated rounds of invasion, replication, and host cell lysis [3]. Following an adequate host immune response, the number of tachyzoites in the host declines, and a subset of tachyzoites differentiate into a quasi-dormant life stage termed the bradyzoite [4]. Bradyzoites, packed within large intracellular tissue cysts, are found predominately in the brain and muscle tissue and are a hallmark of chronic infection, persisting for an indefinite period of time in both rodents and humans [5]. Although advances have been made in understanding how the parasite thrives within host cells and avoids host cell defenses, much remains to be discovered regarding how the unique compartment in which parasites replicate, the parasitophorous vacuole (containing tachyzoites) or the tissue cyst (containing bradyzoites), is optimized and re-modeled throughout the infection process of either life stage.

Towards elucidating the molecular basis behind chronicity, one intensively studied structure specific to the bradyzoite life stage is the cyst wall, which appears as a dense collection of filamentous and vesicular structures underneath a delimiting cyst membrane when viewed by electron microscopy [6]. Many proteins secreted by replicating parasites have been shown to accumulate at the cyst wall, several of which are uniquely expressed in the bradyzoite life-stage [7–9]. Our research group recently identified a set of proteins that were enriched in an *in vitro* derived cyst wall fraction obtained from parasites grown under alkaline-stress conditions [10]. Many of these proteins demonstrated no homology to known proteins of other organisms and were found to be expressed by both tachyzoite and *in vitro* bradyzoite life stages. Although both validated and putative cyst wall proteins lack domains of known function, we were interested in characterizing one putative cyst wall protein that contained a recognizable domain belonging to the PP2C class of serine/threonine phosphatases. We hypothesized that this protein, previously dubbed TgPPM3C [11], could influence the viability of parasites by affecting the phosphorylation status of vacuolar protein substrates.

We tested our hypothesis by first determining the localization of epitope-tagged TgPPM3C in tachyzoite and bradyzoite vacuoles and proceeded with ablating TgPPM3C protein expression. We found that although a minor growth defect was observed in TgPPM3C knockout parasites (ΔTgPPM3C) in human fibroblast cultures *in vitro*, a complete loss of virulence was observed during acute infection in mice, as well as a decrease in brain cyst number during chronic infection. Both phenotypes were rescued by complementation of the TgPPM3C gene. We assessed the phosphoproteome of ΔTgPPM3C cultures compared to cultures of parasites expressing TgPPM3C and found that phosphopeptides from several secreted dense granule proteins were significantly more abundant in ΔTgPPM3C cultures, potentially revealing TgPPM3C substrates. Based on phosphoproteomic and co-immunoprecipitation results, we then assessed whether protein effector export from the parasitophorous vacuole into the host cell was altered in the ΔTgPPM3C strain. We found that the export of two protein effectors, GRA16 and GRA28, were impaired in the ΔTgPPM3C strain, though the export of another effector, IST, was seemingly not compromised when assessed indirectly. We then demonstrated that mutations mimicking the phosphorylation of two GRA16 serine residues phenocopies impaired GRA16 export, suggesting de-phosphorylation is a key process in facilitating the export of GRA16. Hence, our results demonstrate that TgPPM3C is involved in regulating the export of GRA16 and GRA28, and is likely involved in the management of vacuolar protein phosphorylation at large.

## RESULTS

### TgPPM3C is a putative protein phosphatase localized to the lumen of the parasitophorous vacuole

The TgPPM3C gene (Gene ID: TGME49_270320) is predicted to encode a protein with a signal peptide and a catalytic PP2C domain (Fig 1A). Compared to human PPM1A, TgPPM3C contains an extended N-terminal portion relative to the catalytic domain, but still shares conserved amino acid residues involved in phosphate and Mn^2+^/Mg^2+^ binding (Fig 1A, blue and red arrows). To identify the localization of TgPPM3C, the endogenous locus of the gene was epitope-tagged using Cas9 guide RNA targeting the C-terminus of TgPPM3C. A single-copy of the hemagglutinin epitope (HA) was fused to the C-terminus in the Prugniaud Δku80Δhxgprt (PruQ) background. TgPPM3C-HA was shown to partially co-localize with GRA1 in extracellular tachyzoites (Fig 1B), indicating it is likely secreted through the dense granule secretory pathway. Following 36 hours of infection under tachyzoite growth conditions, TgPPM3C localizes to the lumen of the parasitophorous vacuole (Fig 1C). Bradyzoite induction via growth in alkaline serum-depleted media demonstrates TgPPM3C localization to the matrix of *in vitro* tissue cysts 4 days post-infection (Fig 1D), indicating TgPPM3C does not predominately localize to the cyst wall [10].

**Figure 1.**
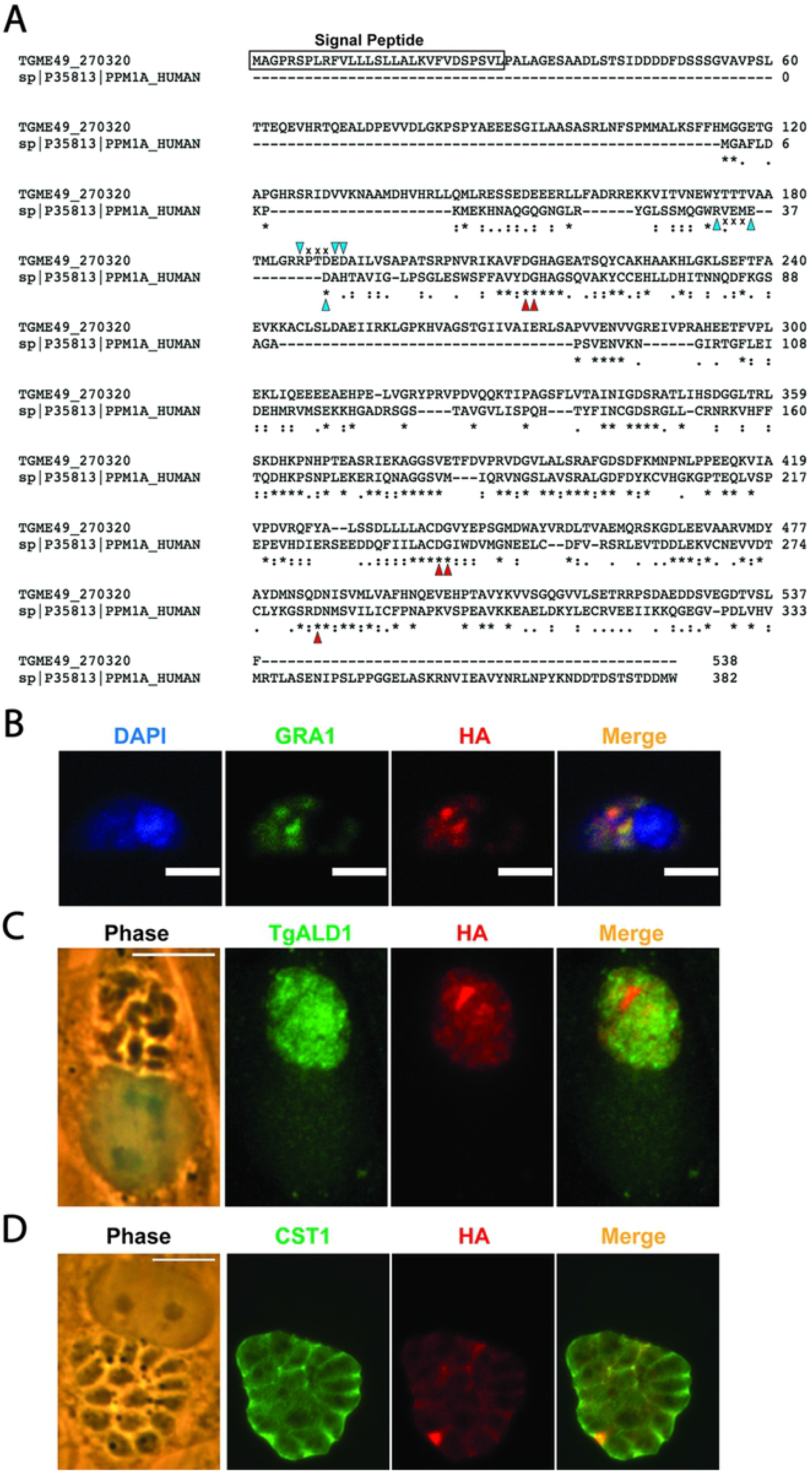
TgPPM3C is a parasitophorous vacuole protein with a PP2C-class phosphatase catalytic domain. **(A)** Amino acid sequence alignment of TgPPM3C (Gene ID: TGME49_270320) and human protein phosphatase 1a (PPM1a), a canonical PP2C class phosphatase, performed with Clustal Omega [53]. TgPPM3C contains a predicted signal peptide (boxed region) and an extended N-terminal domain upstream of the conserved PP2C catalytic domain. Residues important for metal binding are indicated with red arrows. Unaligned but conserved residues important for phosphate and metal binding shared by TgPPM3C and PPM1A are indicated by blue arrows. **(B)** Immunofluorescence image of an extracellular parasite with labeled dense granules (GRA1), endogenously tagged TgPPM3C-HA (HA), and nucleus (DAPI). Partial colocalization is observed between GRA1 and HA, indicating TgPPM3C is likely packaged into in dense granules. **(C)** Immunofluorescence image of a parasitophorous vacuole grown under tachyzoite growth conditions (36 hours post-infection) or bradyzoite growth conditions (**D**, 4 days post-infection). Parasites in **(C)** are labeled with TgALD1, while bradyzoite differentiation in **(D)** is indicated by glycosylated CST1 expression, detected with SalmonE monoclonal antibody. Under both conditions, TgPPM3C is observed within the vacuolar space between parasites, indicating secretion by parasites during intracellular infection.

### ΔTgPPM3C parasites exhibit growth defects *in vitro* and a profound virulence defect *in vivo*

To begin assessing the role of TgPPM3C, expression of TgPPM3C-HA protein was removed by deleting the predicted start codon of TgPPM3C and introducing a multi-stop codon sequence. Using the inserted multi-stop codon sequence as a handle for Cas9 guide RNA targeting, TgPPM3C protein was restored in the ΔTgPPM3C strain to yield a complemented strain (TgPPM3C-COMP). Immunoblotting of tachyzoite lysates demonstrated that TgPPM3C migrates close to the predicted size of the protein (59kDa, Fig 2B). As expected, TgPPM3C-HA protein is completely ablated in the ΔTgPPM3C strain, and comparable levels of TgPPM3C protein are present in the PPM3C-COMP strain compared to the parental TgPPM3C-HA strain (Fig 2B).

**Figure 2.**
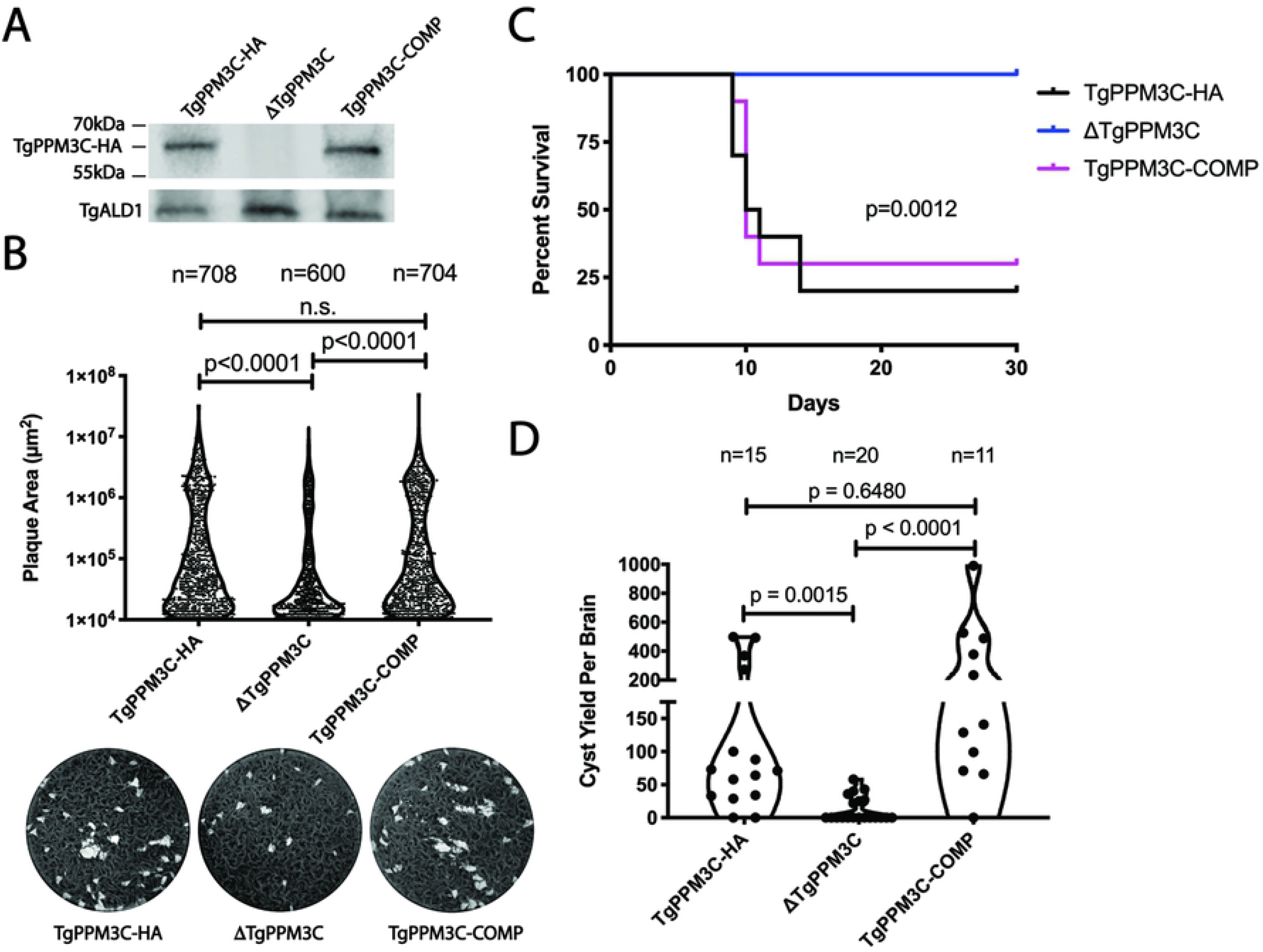
ΔTgPPM3C parasites exhibit growth defects *in vitro* and *in vivo*. **(A)** Immunoblot images obtained from SDS-PAGE separated protein lysates of tachyzoite infected cultures, 24 hours post-infection. Comparable amounts of TgPPM3C protein, migrating close to the predicted size (59kDa), are expressed in TgPPM3C-HA and TgPPM3C-COMP infected cultures but not ΔTgPPM3C infected cultures. TgALD1 was used as a parasite specific loading control. **(B)** Violin plots depicting the distribution of plaque sizes formed by TgPPM3C modified strains after two weeks of growth in human fibroblasts monolayers. Representative images of wells contain plaques from each strain are shown below. 600-700 plaques from each strain, pooled from three independent experiments, were captured and quantified in ImageJ, using the Kruskal-Wallis test to compare each group and generate p-values. A significant decrease in plaque size is observed in the ΔTgPPM3C strain compared to TgPPM3C-HA and TgPPM3C-COMP strains. **(C)** Kaplan-Meier survival curves from C57Bl/6 mice injected intraperitoneally with equal amounts of parasites (16,000) from each strain. 10 mice were injected per group. 20% and 30% of mice survived after 30 days following infection with TgPPM3C-HA and TgPPM3C-COMP parasites respectively, whereas all mice survived following infection with ΔTgPPM3C parasites, indicating ΔTgPPM3C parasites are attenuated. A log-rank test indicates a significant difference in survival curves between groups. Data are representative of two independent experiments. **(D)** Violin plots of cyst burden from individual C57Bl/6 mouse brains, harvested 30 days post-infection following intraperitoneal injection with 16,000 parasites of each strain. Cyst burden is estimated based on cyst counts from one cerebral hemisphere per mouse brain. The numbers of brains from which cysts were quantified are indicated above each plot. Significantly less cysts are formed by ΔTgPPM3C parasites compared to TgPPM3C-HA and TgPPM3C-COMP parasites. Data are pooled from two independent experiments.

To assess for growth defects in the ΔTgPPM3C strain, plaque assays of human foreskin fibroblast monolayers were performed, allowing the TgPPM3C-HA, ΔTgPPM3C, and TgPPM3C-COMP strains to grow for two weeks. The sizes of plaques generated from multiple rounds of the tachyzoite lytic cycle were measured and compared between each strain. A significant decrease in average plaque size was noted in the ΔTgPPM3C strain compared to the TgPPM3C-HA and TgPPM3C-COMP strains (Fig 2C). *In vivo* virulence was assessed next following intraperitoneal infection of C57Bl/6 mice with an inoculum of 16,000 parasites per mouse. Although more than half of the mice infected with TgPPM3C-HA and TgPPM3C-COMP strains succumbed to infection, no mice infected with ΔTgPPM3C parasites perished, revealing a profound defect in virulence (Fig 2D). Cyst burden was measured from the brains of Bl/6 mice chronically infected for 30 days with equal inoculums of each strain. A significant reduction in ΔTgPPM3C cyst number was measured compared to the TgPPM3C-HA and TgPPM3C-strains (Fig 2E). No significant differences in virulence or cyst burden were observed between the TgPPM3C-HA and TgPPM3C-COMP strains (Fig 2D-E).

### Co-immunoprecipitation of TgPPM3C-HA reproducibly enriches only a few parasite proteins

Based on the PP2C domain of TgPPM3C, we hypothesized that secreted vacuolar proteins normally dephosphorylated by TgPPM3C remain phosphorylated in ΔTgPPM3C parasites. As a consequence, the loss of phospho-regulation of one or several key vacuolar proteins results in the virulence and growth defects observed in the ΔTgPPM3C strain. We inspected tachyzoite vacuoles from TgPPM3C-HA and ΔTgPPM3C strains by transmission electron microscopy and found no differences in gross morphology (Figure S1), demonstrating that obvious disruption of the *Toxoplasma* parasitophorous vacuole architecture was not responsible for the ΔTgPPM3C phenotype. To identify TgPPM3C interacting parasite proteins, we performed co-immunoprecipitation experiments from tachyzoite infected cultures 24 hours post-infection, using anti-HA magnetic beads to pulldown TgPPM3C-HA protein. As a negative control, Co-IPs of non-epitope tagged PruQ parasites with anti-HA magnetic beads were also performed. Based on two independent experiments, apart from TgPPM3C-HA, we identified two GRA proteins (GRA7, GRA9) and two proteins known to be secreted into the parasitophorous vacuole (MAG1, MYR1) as significantly enriched in both experiments (p < 0.05, log2 fold-change > 1.5) (Table 1, Dataset S1). Intriguingly, the MYR1 protein is known to be involved in the export of parasite effectors from within the parasitophorous vacuole into the host cell across the parasitophorous vacuole membrane [12]. Nonetheless, despite the reproducible enrichment of a few proteins known to be secreted into the parasitophorous vacuole, it is possible that other substrates of TgPPM3C were not identified by the Co-IP approach, particularly if de-phosphorylation events occur rapidly with transient interactions between TgPPM3C and protein substrates.

**Table 1.**
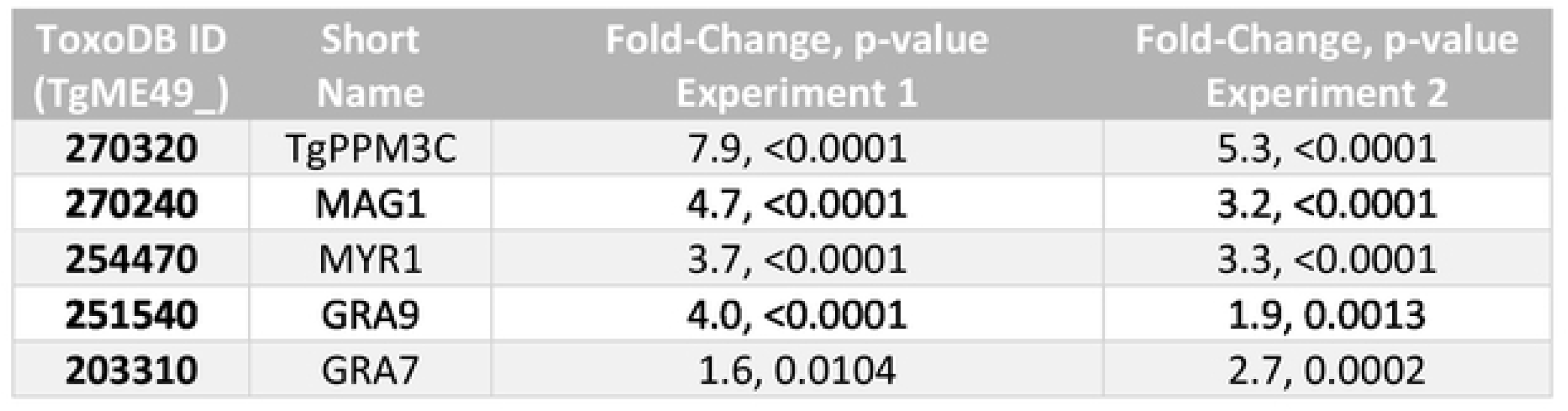
Five parasite secreted proteins are significantly enriched by TgPPM3C-HA Co-IP. Two independent Co-IP experiments were performed to identify TgPPM3C interacting proteins. Fold change over eluates from untagged control samples and p-values from each experiment are provided. Only protein hits with greater than 1.5 fold-change over control and p-value < 0.05 in both experiments are shown.

### Phosphoproteomic comparison ofΔTgPPM3C and TgPPM3C-HA parasite cultures reveal putative phosphoprotein substrates

PP2C-class phosphatases identified in various organisms thus far exhibit no consensus target motif that allows for *a priori* PP2C substrate prediction. Therefore, we turned to a label-free phosphoproteomic approach to gain insights on putative TgPPM3C substrates in an unbiased fashion. Human foreskin fibroblast monolayers grown in 15cm diameter culture dishes were heavily infected in triplicate at a multiplicity of infection of five (MOI 5) with either TgPPM3C-HA (referred to as wild type, or “WT”, for the purposes of this experiment,) or ΔTgPPM3C parasites and allowed to grow under tachyzoite growth conditions for 36 hours, allowing for large vacuoles to be obtained prior to parasite egress. Protein was harvested from infected cultures using 5% SDS lysis buffer and digested using S-Trap (Protifi) columns. Phosphopeptides were subsequently enriched with titanium dioxide beads and processed through LC-MS/MS to obtain peptide spectra. To normalize phosphopeptide abundance by protein abundance, the flow-through fraction from titanium dioxide bead enrichment (i.e. non-phosphorylated proteins) were collected from each replicate and also analyzed by LC-MS/MS.

Analysis of LC-MS/MS spectra revealed a robust, high confidence detection of 5,882 phosphopeptides from both host cell and parasite (Dataset S2). Among the 2,388 unique parasite phosphopeptides identified, 128 phosphopeptides were found to have significantly altered abundance in the ΔTgPPM3C strain, as defined by a p < 0.05 cut-off and log_2_ fold-change of less than −1.2 or greater than 1.2 (Fig 3A, red points). Of these, 118 total phosphopeptides were significantly more abundant in the ΔTgPPM3C strain compared to the parental strain, whereas only 10 phosphopeptides, largely from hypothetical proteins of unknown function, were significantly less abundant (Fig 3A). Interestingly, more than half (79) of all differentially abundant phosphopeptides belong to proteins previously known to be secreted into the lumen of the parasitophorous vacuole (Dataset S2), suggesting TgPPM3C predominately affects the phosphorylation status of vacuolar proteins. Hypergeometric testing was performed to identify parasites compartments that were significantly enriched by all detected phosphopeptides from both wild-type and ΔTgPPM3C samples, as compared to the entire *Toxoplasma* proteome (Figure 3C, upper panel). Enrichment could be seen for several parasite compartments, including compartments containing non-secreted proteins such as the nucleus and cytoplasm (Figure 3C, upper panel). In contrast, among the phosphopeptides significantly more abundant in ΔTgPPM3C samples, dense granule, rhoptry, and micronemes were the only significantly enriched compartments based on hypergeometric testing (Figure 3D, upper panel). The most robust compartment enrichment observed for this analysis was the dense granule, suggesting that the majority of phosphoproteins significantly affected by the absence of TgPPM3C are dense granule proteins secreted into the parasitophorous vacuole (Figure 3D, upper panel). Among the phosphopeptides that were more significantly more abundant from ΔTgPPM3C samples, no amino acid motif appeared to be enriched when inspected within a −15 and +15 amino acid window (Figure 3D, lower panel) as compared to −15 and +15 amino acid window analysis of randomly selected *Toxoplasma* phosphopeptides (Figure 3C, lower panel). This result is in agreement with the notion that PP2C phosphatases do not have clear amino acid target motifs that are recognized and dephosphorylated.

**Figure 3.**
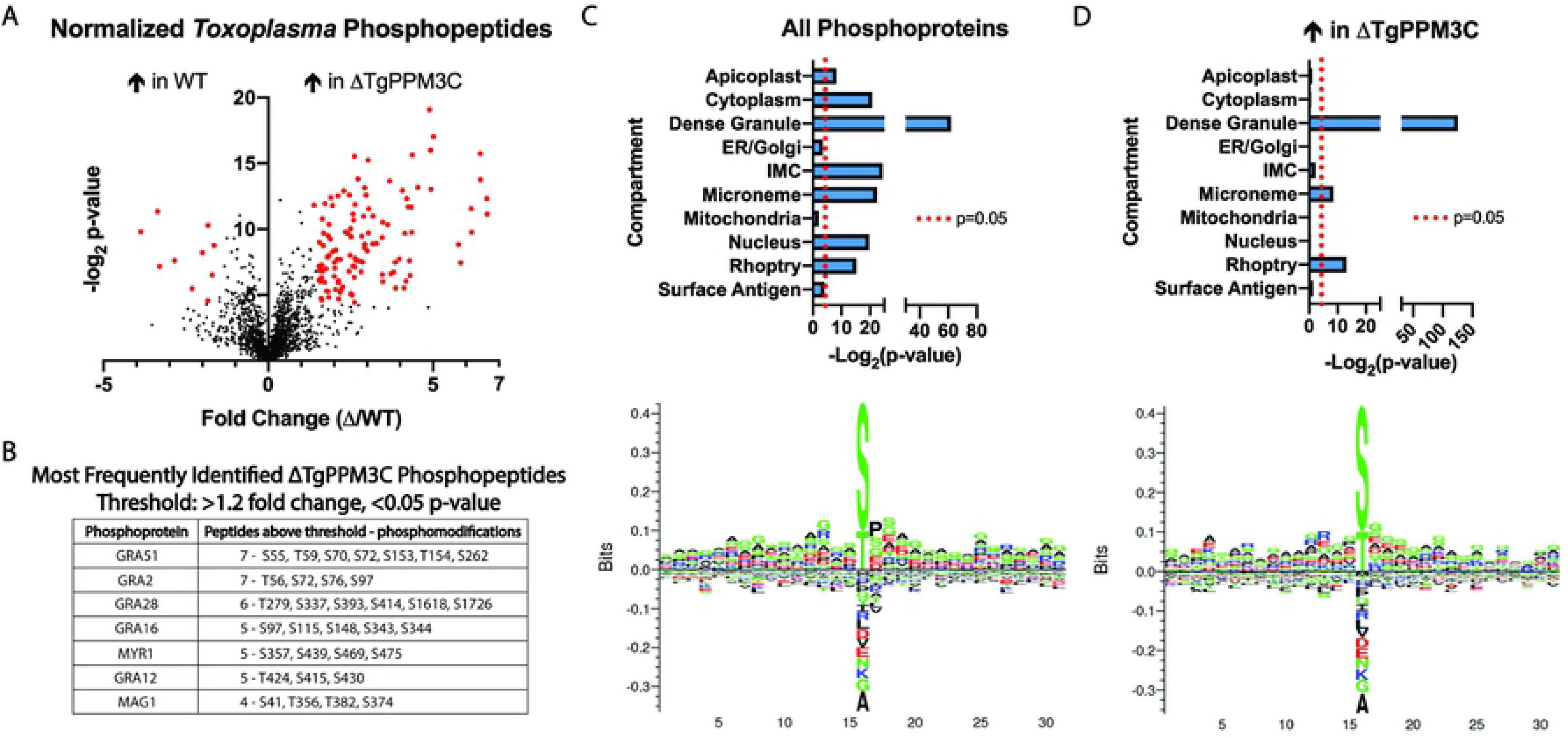
Phosphoproteomic analysis of TgPPM3C-HA (WT) and ΔTgPPM3C parasites identifies phosphopeptides that are more abundant in ΔTgPPM3C cultures. **(A)** Volcano plot depicting significant fold-changes in *Toxoplasma* phosphopeptides from ΔTgPPM3C samples over TgPPM3C-HA (WT) samples. Phosphopeptides with fold changes greater than 1.2 or less than −1.2 and with −log_2_ p-values greater than 4.32 (equivalent to p < 0.05) are highlighted in red. Using this criteria, 118 phosphopeptides are more abundant in ΔTgPPM3C cultures, as opposed to 10 phosphopeptides more abundant in WT cultures, suggesting that the absence of TgPPM3C predominately results in the accumulation of phosphoproteins that are normally dephosphorylated by TgPPM3C. Phosphopeptides are normalized to their respective protein abundance, detected in the flow-through fraction of titanium dioxide beads used for phosphopeptide enrichment. **(B)** A table listing phosphoproteins detected in this dataset containing the most phosphopeptides above a 1.2 fold change and p < 0.05 threshold. Notably, two exported protein effector proteins (GRA16 and GRA28) and a protein involved in facilitating effector export (MYR1) are among these proteins. **(C)** Top – Results from hypergeometric testing to identify significant enrichment for subcellular parasite compartments among all phosphopeptides detected from wild-type and ΔTgPPM3C samples, compared to the *Toxoplasma* proteome. Various compartments are significantly enriched. Bottom – Amino acid motif, generated by Seq2Logo [52], derived from randomly selected phosphopeptides identified in wild-type and ΔTgPPM3C samples using a −15 and + 15 amino acid window with respect to the phosphoserine or phosphothreonine. As expected, no clear consensus motifs are evident. **(D)** As described in **(C)**, except that hypergeometric testing (top) and amino acid motif generation (bottom) was performed solely with phosphopeptides identified as significantly enriched in ΔTgPPM3C samples. A clear enrichment for the dense granule compartment and to a lesser extent rhoptry and microneme compartments are observed among these phosphopeptides, although no consensus amino acid motif is evident.

### GRA16 and GRA28 export from the parasitophorous vacuole is impaired in ΔTgPPM3C parasites

In line with our initial hypothesis, we focused our studies on phosphoproteins identified by LC-MS/MS with phosphopeptides that were significantly more abundant in ΔTgPPM3C cultures. Intriguingly, this list was well-represented by phosphopeptides belonging to two proteins known to be exported beyond the parasitophorous vacuole into the host cell nucleus, GRA16 [13] and GRA28 [14], as well as the afore-mentioned MYR1 protein (Fig 3B). Based on these observations, and the detection of MYR1 from TgPPM3C Co-IP experiments (Table 1), we tested whether defects in effector export were evident in the ΔTgPPM3C strain, which might at least partially explain the observed growth and virulence defects, since parasites lacking MYR1 or GRA16 exhibit a profound loss of virulence in mice [12, 13]. Transient transfections with plasmids encoding epitope tagged GRA16 and GRA28 under control of their endogenous promoters were performed with PruQ and ΔTgPPM3C parasites, and the host nuclear accumulation of each effector was assessed 24 hours post-infection. A significant reduction in GRA16 and GRA28 nuclear accumulation was observed during infection with the ΔTgPPM3C strain as compared to the PruQ strain, likely indicating defects in effector export (Fig 4A).

**Figure 4.**
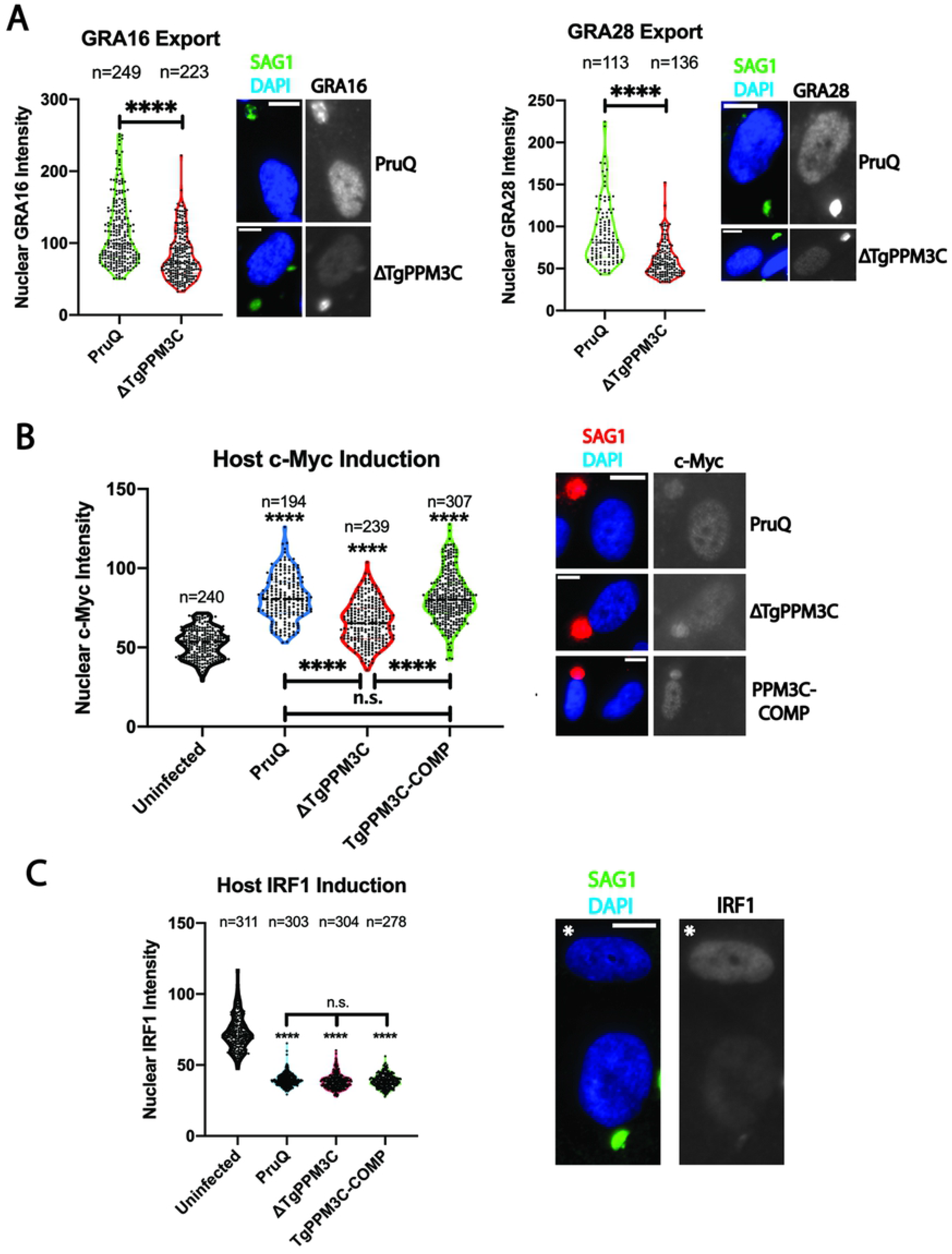
ΔTgPPM3C parasites exhibit partial defects in protein effector export. **(A)** Violin plots of GRA16-3xHA and GRA28-3xHA accumulation in host nuclei infected with either PruQ or ΔTgPPM3C parasites, transiently transfected with epitope tagged effector constructs. GRA16 and GRA28 fluorescence were quantified from fibroblast nuclei containing a single HA-positive parasite vacuole. A significant decrease in GRA16 and GRA28 host nuclear accumulation is observed during ΔTgPPM3C infection, indicating defects in effector export from the parasitophorous vacuole. Data were collected from three technical replicates. Representative images from which effector intensity were quantified are shown on the right. Antibody to SAG1 was used as a parasite marker, DAPI as a host nucleus marker, and anti-HA antibody was used to detect GRA16-3xHA and GRA28-3xHA effectors. **(B)** Violin plots of c-Myc induction in host nuclei infected with either PruQ, ΔTgPPM3C, or TgPPM3C-COMP parasites. Host c-Myc fluorescence was quantified from fibroblast nuclei containing a single parasite vacuole. A significant decrease in host c-Myc induction is observed during ΔTgPPM3C infection compared to PruQ and TgPPM3C-COMP infections. Notably, significant induction of host c-Myc expression over uninfected host cells are observed during infection with all three strains (asterisks above each plot). Data are pooled from three independent experiments using different batches of host cells and parasites. Representative images from which c-Myc intensity were quantified are shown on the right. Antibody to SAG1 was used as a parasite marker and DAPI as a host nucleus marker. **(C)** Violin plots of IRF1 induction in host nuclei infected with either PruQ, ΔTgPPM3C, or TgPPM3C-COMP parasites and stimulated with IFN-γ for six hours prior to fixation. Host IRF1 fluorescence was quantified from fibroblast nuclei containing a single parasite vacuole. Although each strain significantly attenuated IRF1 induction compared to uninfected cells (asterisks), no significant differences in the extent of IRF1 suppression are observed between any of the strains. Data are pooled from three independent experiments using different batches of host cells and parasites. A representative image from a PruQ infected fibroblast and uninfected fibroblast (white asterisk) is shown underneath the violin plot. Antibody to SAG1 was used as a parasite marker and DAPI as a host nucleus marker. For all violin plots in this figure, **** asterisks indicate p < 0.0001 and n.s. indicates no significant difference. The number of observations made for quantification are provided above each violin plot. Kruskal-Wallis tests were performed to calculate p-values.

HFF monolayers were next infected with either PruQ, ΔTgPPM3C, or TgPPM3C-COMP parasites and assayed for alterations in host c-Myc upregulation, which is known to be dependent on MYR1 and GRA16 export [12, 15]. The results demonstrated a significant reduction in host c-Myc upregulation during infection with ΔTgPPM3C parasites compared to the PruQ and TgPPM3C-COMP strains (Fig 4B). Notably, all strains significantly induced host c-Myc upregulation compared to uninfected serum-starved fibroblasts, indicating that the absence of TgPPM3C does not result in a complete loss of effector translocation, in agreement with recent observations made by colleagues on ΔTgPPM3C mutants in a different parasite strain [16].

To test whether effector export defects were specific to GRA16 and GRA28 or applied more broadly to MYR1 dependent effectors, we also assayed for defects in the export of parasite effector TgIST indirectly by querying alterations in host IRF1 upregulation, which is known to be inhibited in a TgIST-dependent fashion in infected cells stimulated with IFN-γ [17, 18]. The results showed no significant differences in host IRF1 attenuation following IFN-γ stimulation when comparing PruQ, ΔTgPPM3C, and TgPPM3C-COMP infections (Fig 4C), suggesting that TgIST effector export is not impaired in the ΔTgPPM3C strain.

### Phosphomimetic mutations of GRA16 recapitulate vacuolar export defects

Based on the findings obtained from host IRF1 attenuation, the defects observed in the ΔTgPPM3C strain suggest that GRA16 and GRA28 export are specifically impaired due to their altered phosphorylation status, as opposed to impaired function of MYR1. To test this hypothesis for the GRA16 effector, we mutagenized two serine residues in the GRA16 protein (S97 and S148) whose phosphorylation was robustly detected in ΔTgPPM3C parasites and absent in TgPPM3C-HA parasites (Fig 3B, Supplementary Dataset 1). These residues fall within the R1 and R2 tandem repeat regions (R1: 59-100aa, R2: 125-168aa) previously shown to be critical for GRA16 export [13, 19]. Phosphomimetic (S97E/S148E) and phosphoablative mutations (S97P/S148P) were introduced into constructs encoding epitope-tagged GRA16 under control of the endogenous promoter (Fig 5A). For phosphoablative mutations, proline substitutions were used instead of alanine so as to maintain protein intrinsic disorder within the mutant GRA16 sequence, which is known to be another critical feature that facilitates GRA16 export [19]. Proline, unlike alanine, promotes protein disorder and is commonly found within intrinsically disordered proteins [20]. Following transfection of PruQ parasites, host nuclear GRA16 accumulation was assessed by immunofluorescence 24 hours post-infection with polyclonal parasites expressing either wild-type (unmodified), phosphomimetic, or phosphoablative GRA16 constructs. The results demonstrated that both mutant GRA16 proteins were significantly less abundant within infected host cell nuclei compared to the wild type GRA16 protein, as measured by immunofluorescence intensity (Fig 5B). However, th phosphomimetic (S97E/S148E) GRA16 was found to be significantly less abundant in host nuclei compared to phosphoablative (S97P/S148P) GRA16 (Fig 5B), indicating that the glutamate substitutions compromised GRA16 export more drastically than the proline substitutions. Although we cannot completely exclude the possibility that the mutant GRA16 proteins might somehow be less stable in the host nucleus compartment compared to wild type GRA16 protein, we believe this possibility is less likely, and that the findings most likely reflect a decreased export efficiency from the parasitophorous vacuole.

**Figure 5.**
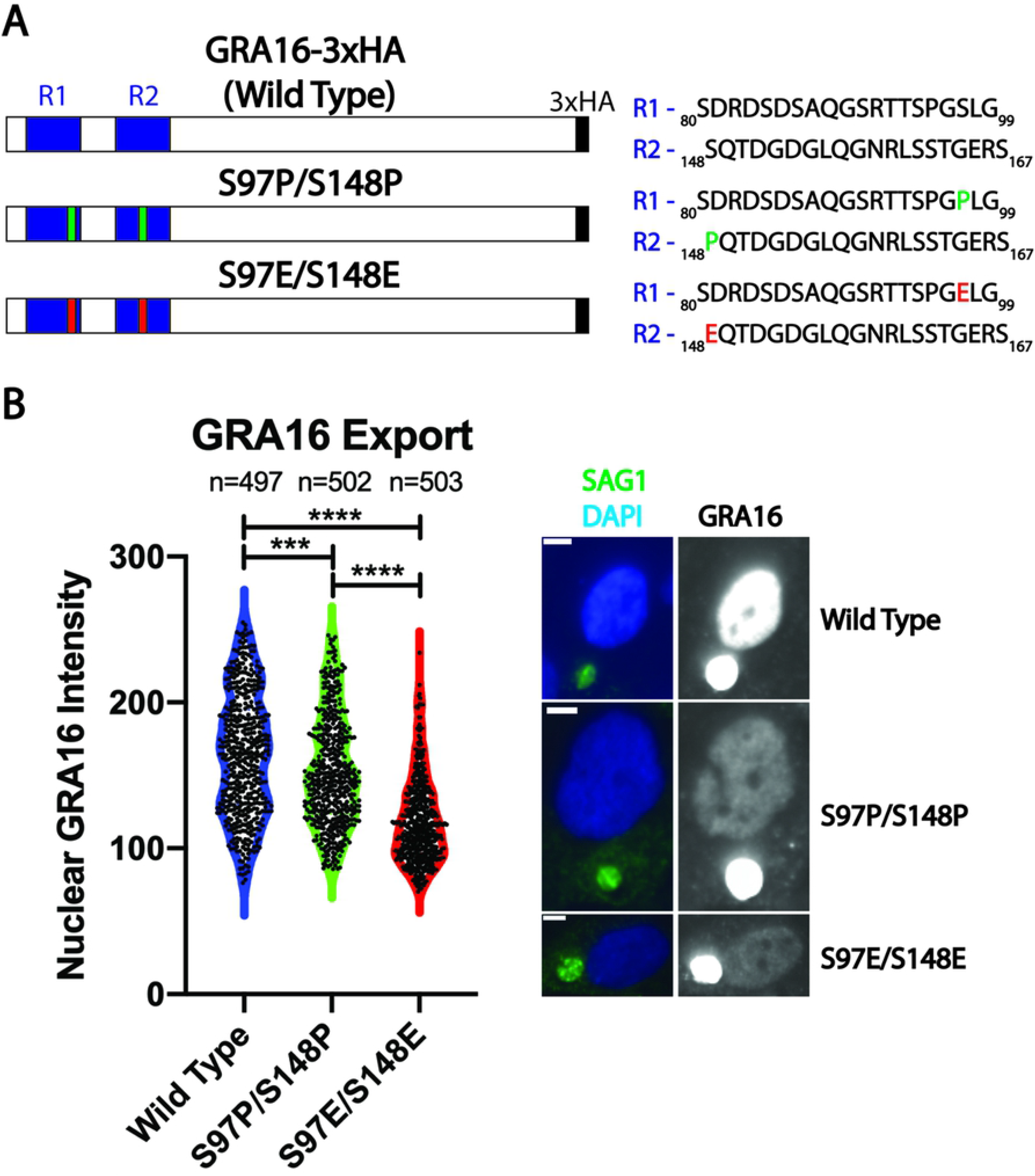
Phosphomimetic mutations impair the export of GRA16 from the parasitophorous vacuole. **(A)** Schematic of GRA16-3xHA constructs, including the partial sequence and location of previously described tandem repeat regions (R1 and R2). Mutations were designed to introduce either proline (S97P/S148P) or glutamate (S97E/S148E) in place of serine 97 and serine 148. **(B)** Violin plots and representative images of GRA16 intensity in host nuclei infected with polyclonal parasites expressing either wild type, S97P/S148P, or S97E/S148E GRA16-3xHA constructs. Nuclear GRA16 intensity was quantified from fibroblast nuclei containing a single HA-positive parasite vacuole. A significant decrease in nuclear S97P/S148P and S97E/S148E GRA16 intensity is observed compared to wild type GRA16 intensity, while S97E/S148E GRA16 nuclear intensity values was also found to be significantly less intense compared to values recorded from S97P/S148P infections. Data are pooled from three independent experiments using different batches of host cells and parasites. Representative images from which GRA16 intensities were quantified are shown on the right. Antibody to SAG1 was used as a parasite marker and DAPI as a host nucleus marker. *** asterisks indicate p < 0.001, **** indicates p < 0.0001. The number of observations made for quantification are provided above each violin plot. Kruskal-Wallis tests were performed to calculate p-values.

## DISCUSSION

PP2C-class serine/threonine protein phosphatases are widespread throughout all domains of life. Through the regulation of protein phosphorylation status, PP2C phosphatases influence cellular activity and signaling in diverse fashions, ranging from growth to survival [21]. An expansion of genes encoding PP2C catalytic domains is noteworthy in plants, some of which play major roles in stress-response signaling pathways [22]. Intriguingly, *Toxoplasma* exhibits several plant-like features, such as the expression of calcium dependent protein kinases [23], plant-like AP2 transcription factors [24], as well as the expansion of genes predicted to encode PP2C-class phosphatases [11]. In good agreement with this observation is a previous study on phosphatase activity in parasite lysates, which demonstrated that PP2C was by far the more prevalent phosphatase activity compared to PP2A activity, as determined through the use of phospho-casein as a substrate and PP2A-class phosphatase inhibitors [25]. TgPPM3C is one of several PP2C genes predicted to encode signal peptides, suggesting several PP2C phosphatases may enter the secretory pathway and be packaged into secretory organelles. Indeed, a previous study demonstrated that one of these PP2C phosphatases, dubbed PP2C-hn, is packaged into rhoptry organelles and secreted into the host cell nucleus at the time of invasion [26]. In contrast, we show that TgPPM3C is at least partially packaged into dense granules (Fig 1B), which are organelles that continuously release their contents into the parasitophorous vacuole during intracellular infection [27]. Transcriptomic data deposited on ToxoDB [28] demonstrate TgPPM3C mRNA transcripts in tachyzoites, bradyzoites, and merozoites, suggesting a shared role for this phosphatase during different *Toxoplasma* life stages. Both the catalytic domain and N-terminal extension of TgPPM3C appears to be largely conserved across three strains representative of three major lineages of *Toxoplasma* (GT1, ME49, and VEG; Fig S2). Furthermore, a high degree of TgPPM3C homology is evident in *Neospora caninum* and *Hammondia hommondi* (Fig S3), possibly indicating a common need for this particular PP2C phosphatase among the close relatives of *Toxoplasma*.

The conserved PP2C catalytic domain forms a tertiary structure in which a beta-sandwich surrounded by alpha helices forms a pocket that allows metal-coordinating and phosphate binding residues to catalyze de-phosphorylation [29] (Fig 1A). In agreement with homology to the PP2C domain, the X-ray crystal structure of the TgPPM3C catalytic domain obtained by Almo and colleagues [30] demonstrates that the catalytic domain of TgPPM3C appears to adopt the appropriate tertiary structure to mediate de-phosphorylation (Fig 6A-B). A largely similar catalytic pocket architecture can be observed in TgPPM3C (Fig 6A-B) compared to human PPM1A (Fig 6C-D). Notably, although the TgPPM3C active site is putative, this region forms a robust acidic pocket that likely attracts Mg^2+^/Mn^2+^ cations (Fig 6B, bottom panel), as observed in human PPM1A (Fig 6D, bottom panel). Although a phosphate ion can be seen in the PPM1A crystal structure (Fig 6C-D), the sulfate ions used in conditions to crystallize TgPPM3C were not found to reside in the active site pocket. The TgPPM3C crystal analyzed here also does not contain the N-terminal extension domain of TgPPM3C, which conceivably could be involved in determining substrate preference and/or regulating catalytic activity. The flap domain (green regions in Fig 6) is a common feature among PP2C phosphatases that is thought to be involved in substrate preference by docking phosphopeptides along the groove formed between the active site and flap domain [29]. Indeed, this configuration has been demonstrated for the *Arabidopsis* phosphatase TAP38/PPH1, which demonstrates a clear preference for a phosphopeptide from LHCII by virtue of basic residues adjacent to the phosphothreonine substrate that mediate electrostatically favorable interactions with the acidic TAP38/PPH1 catalytic pocket [31]. TgPPM3C contains an additional cleft not observed in human PPM1A beside the flap domain and putative active site (Fig 6A-B, white region); this groove may also be involved in phosphoprotein substrate docking, although more experiments are needed to test this hypothesis.

**Figure 6.**
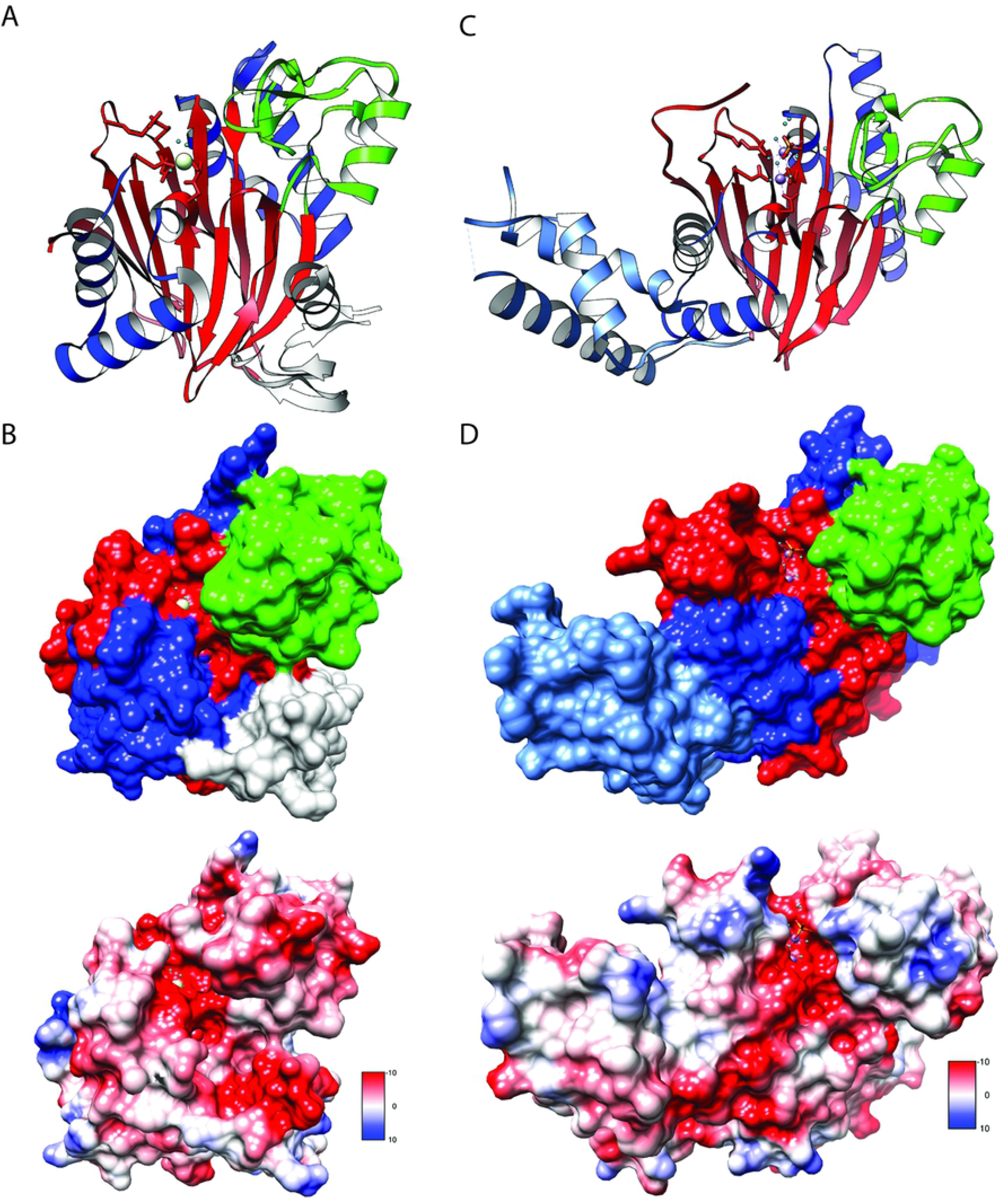
The X-ray crystal structure of TgPPM3C catalytic domain demonstrates a largely conserved tertiary structure. **(A)** Ribbon projection of TgPPM3C crystal structure. The characteristic beta sandwich (red) formed by adjacent beta sheets can be seen coordinating a Praseodymium cation (Pr^3+^) (white circle) in the putative active site of the enzyme. The side chains of the conserved metal and phosphate coordinating amino acids are shown, along with water molecules (small cyan circles) in this region. Alpha helices (blue) surround the beta sandwich. The flap domain (green) is seen above the beta-sandwich pocket, along with an additional domain (white) present beneath the flap domain, which is not seen in the human PPM1A crystal structure. **(B)** Top – surface mesh rendering of TgPPM3C, demonstrating the likely catalytic pocket in which the Pr^3+^ ion is found. Coloring scheme is as described in **A.** Bottom – calculated Coulombic potential for the TgPPM3C surface mesh, demonstrating a strongly acidic (red) putative catalytic pocket with no basic residues (blue) near the beta-sandwich cleft. **(C)** Ribbon projection of the human PPM1A crystal structure. The beta sandwich (red) can be seen coordinating two Mg^2+^ cations (purple circles) in the active site, along with a phosphate ion (orange stick figure) just above the Mg^2+^ cations. The side chains of the conserved metal and phosphate coordinating amino acids are shown, along with water molecules (small cyan circles) in this region. The flap domain is shown in green, and the C-terminal domain extending away from the center of the enzyme is shown in light blue. **(D)** Top – surface mesh rendering of PPM1A, demonstrating a similar active site pocket as seen in TgPPM3C. Coloring scheme is as described in **C.** Bottom – calculated Coulombic potential for the PPM1A surface mesh, demonstrating the characteristic acidic (red) catalytic pocket and a few basic residues (blue) above the catalytic pocket. Molecular graphics and analyses of crystal structures were prepared using UCSF Chimera [54] and the publicly available structures of the TgPPM3C catalytic domain (Protein Data Bank ID: 2ISN) and human PPM1A (Protein Data Bank ID: 1A6Q).

Current data on PP2C substrate recognition suggest that no specific consensus motifs are recognized by the catalytic domain; rather, it is thought that interacting protein partners and/or divergent targeting domains present in PP2C phosphatases direct the enzyme to substrates [32]. We speculate that interacting proteins such as MYR1 and/or the N-terminal extension of TgPPM3C may serve as the entities responsible for targeting this phosphatase to the appropriate substrates. Along these lines, we observed the lack of a consensus motif among the phosphopeptides that were more abundant in ΔTgPPM3C cultures (Fig 3D, lower panel), which we propose largely represent direct TgPPM3C phosphoprotein substrates. Support for this notion is provided by the fact that more than half of these phosphopeptides belong to proteins known to be secreted into the parasitophorous vacuole, either in soluble or membrane bound forms, (Fig 3A, Dataset S2), which is also where TgPPM3C is located during infection. Furthermore, the dense granule compartment demonstrated the most significant enrichment based on hypergeometric testing of phosphopeptides that were significantly more abundant in ΔTgPPM3C cultures (Fig 3D, upper panel), although we recognize that a bias toward the enrichment of dense granule proteins is evident in our phosphoprotein dataset based on hypergeometric testing of all detected phosphopeptides (Fig 3C, upper panel). While their localization has been described, the specific functions of many GRA/vacuolar proteins are poorly understood and remain to be fully characterized.

Protein phosphorylation within the parasitophorous vacuole has been previously described for GRA7 [33], and several other proteins have been suggested as potentially phosphorylated post-secretion into the vacuole [33, 34]. Putative vacuolar kinases have been previously described, such as ROP21 and ROP27, which were recently shown to localize to the parasitophorous vacuole lumen and cyst matrix of tachyzoite vacuoles and *in vitro* tissue cysts respectively, but not to rhoptry organelles [35]. The simultaneous deletion of ROP21 and ROP27 results in reduced cyst burdens in mouse brain, suggesting that their catalytic activity within the cyst matrix is pivotal to the establishment of cysts during chronic infection [35].

Measures to counterbalance vacuolar protein phosphorylation are likely provided by vacuolar phosphatases. Interestingly, among the several GRA7 phosphopeptides identified in our dataset, one phosphopeptide was significantly less abundant in ΔTgPPM3C cultures (T90, Dataset S2), indicating TgPPM3C may affect the phosphorylation status of proteins indirectly through the regulation of kinases and/or other phosphatases. Indeed, our phosphoproteomic dataset also identified a significant increase in one phosphopeptide from another putative PP2C class phosphatase, PPM3A (S167, Dataset S2), predicted by LOPIT data on ToxoDB to localize to dense granules [28], indicating potential PPM3A secretion into the vacuole and cross-talk between PP2C phosphatases. The vacuolar kinase WNG1 was recently shown to be pivotal in organizing the intravacuolar network (IVN) within the parasitophorous vacuole through the phosphorylation of GRA proteins, in turn affecting the affinity of some GRA proteins to lipid membranes [36]. Phosphopeptides from GRA2 were frequently identified as more abundant in ΔTgPPM3C cultures, including peptides with phosphosites identified as down-regulated in ΔWNG1 cultures from the afore-mentioned study [36] (T56 and S72, Figure 3B). Although we observed no notable differences in the IVN of vacuoles formed by the ΔTgPPM3C strain (Fig S1), we cannot exclude the possibility that abnormalities in ΔTgPPM3C parasitophorous vacuoles may be more evident in the tissue cysts containing bradyzoites or at different time points during tachyzoite development. The regulation and consequences of protein phosphorylation and de-phosphorylation specifically within the parasitophorous vacuole will undoubtedly require extensive investigation, as much remains to be understood regarding the functional outcome of phosphorylation detected in various vacuolar proteins.

The export of protein effectors from the *Toxoplasma* parasitophorous vacuole is a fast-developing area of inquiry [37]. Exported effectors such as GRA16 and TgIST have been elegantly characterized and shown to influence host cell signaling in profound ways that benefit parasite intracellular development [13, 17, 18]. Several effectors exported for the parasitophorous vacuole have been discovered apart from GRA16, GRA28, and TgIST [38–41], while the list of proteins necessary for the export of these proteins has also been growing [16, 42–44], including an acid phosphatase domain containing protein, GRA44. Our data demonstrate that although TgPPM3C is not essential for effector export, this phosphatase likely optimizes effector export based on the deficiencies observed in GRA16 and GRA28 export and host c-Myc induction (Fig 4A-B). TgPPM3C does not appear to influence the export of all effectors, as no defects were seen in the downstream silencing of IFN-γ mediated by TgIST export (Fig 4C). Indeed, we also directly tested whether TgIST export is compromised via transient transfection of PruQ and ΔTgPPM3C strains with a TgIST-3xHA encoding construct and found no obvious differences in the host nuclear accumulation of this effector (data not shown). Hence, the altered phosphorylation status of MYR1 detected in ΔTgPPM3C cultures does not appear to significantly influence the export of all MYR1 dependent effectors. We speculate that the MYR1 phospho-modifications that are more abundant in the ΔTgPPM3C strain may be involved in contacts with only select effector proteins, such as GRA16 and GRA28, and/or potentially other non-effector proteins.

If effector protein dephosphorylation is necessary for efficient export across the parasitophorous vacuole membrane, it is unclear what purpose phosphorylation serves prior to export. All effector proteins exported from the parasitophorous vacuole thus far are predicted to exhibit a large degree of protein intrinsic disorder; a property that is highly correlated with the propensity of proteins to aggregate and form a phase separated state [45]. Perhaps by introducing negatively charged regions that repel each other, vacuolar phosphorylation of intrinsically disordered effector proteins prevents their aggregation. Precedence for this notion can be found in the P granules of *C. elegans* embryos, in which phosphorylation of intrinsically disordered proteins drives de-formation of phase separated granules whereas phosphatase activity induces the opposite effect [46], although an obvious difference in this system is that P granule formation involves RNA as a seed for intrinsically disordered proteins with RNA binding motifs to form a phase separated state. The GRA45 protein was recently shown to facilitate effector protein export by seemingly preventing the aggregation of vacuolar GRA16 and GRA24 [44], which provides support for the notion that effector proteins can aggregate within the parasitophorous vacuole, potentially due to their predicted disordered state. Based on the impaired export of phosphomimetic GRA16, we propose that the de-phosphorylation of GRA16 and GRA28 somehow enables these effectors to traverse the hypothetical vacuolar protein translocon more efficiently. Intriguingly, in RS-repeat rich proteins, the phosphorylation of multiple serine residues has been shown to induce a more rigid and less disordered protein structure [47]. Given that the export of phosphoablative GRA16 was also impaired, and that proline substitutions should in principle maintain protein intrinsic disorder, we speculate that certain non-phosphorylated serine motifs are preferentially recognized by the vacuolar translocon, as opposed to a model where the dephosphorylation of serine serves as a switch from an ordered-to-disordered protein state. Experiments probing the tertiary structure (or lack thereof) of GRA16 are clearly needed to fully understand the structural consequences of phosphorylation in this effector protein, particularly within the R1 and R2 tandem repeat regions.

Altogether, we propose a model whereby TgPPM3C, following secretion through dense granules into the parasitophorous vacuole, de-phosphorylates several vacuolar phosphoprotein substrates including GRA16 and GRA28, which allows these effectors to be efficiently exported and proceed with host cell manipulation, thus optimizing parasite growth and virulence. To validate this model, clearly more investigations are needed on TgPPM3C catalytic activity and substrate preference, where TgPPM3C activity is detected *in situ*, and what other consequences arise from the lack of dephosphorylation in vacuolar proteins not explored in this study.

## MATERIALS AND METHODS

### Cell Culture

PruΔku80Δhxgprt LDH2-sfGFP parasites [48] were continuously passaged in human foreskin fibroblasts (HFF:ATCC:CRL-1634; Hs27) in a 37°C, 5% CO_2_ incubator using Dulbecco’s Modified Eagle Media (DMEM, Gibco) supplemented with 10% fetal calf serum, 1% L-glutamine, and 1% penicillin and streptomycin. Cultures were regularly inspected and tested negative for mycoplasma contamination. Bradyzoite induction was performed at the time of invasion by replacing growth media with bradyzoite induction media (50 mM Hepes, pH 8.2, DMEM supplemented with 1% FBS, penicillin and streptomycin) prior to infection of HFFs with egressed tachyzoites. Bradyzoite induced cultures were maintained in a 37°C incubator without CO_2_, with induction media replaced every 2 days.

### Cloning and Parasite Transfections

For a full list of primers used for cloning and genetic manipulations, refer to Table S1. Briefly, for TgPPM3C epitope tagging, a single guide RNAs (sgRNA) targeting the C-terminus of the TgPPM3C gene was cloned into the p-HXGPRT-Cas9-GFP plasmid backbone using KLD reactions (New England Biolabs), as previously described [49]. Donor sequences for homology mediated recombination were generated by amplifying a 1xHA tag, the 3’UTR of HXGPRT, and a DHFR mini-cassette to confer pyrimethamine resistance from the previously described pLIC-3xHA-DHFR plasmid backbone [10]. Primers used to amplify this donor sequence contained overhangs with 40bp homology to the C-terminus and 3’UTR, as well as mutations in the Cas9 target site to prevent re-targeting by Cas9. To knockout the TgPPM3C gene (and generate ΔTgPPM3C strains), a sgRNA targeting the N-terminus of TgPPM3C was designed and cloned into the afore-mentioned Cas9 plasmid backbone. Donor sequences containing a multi-stop codon sequence and lacking the TgPPM3C start codon were designed for co-transfection with the TgPPM3C knockout sgRNA. To reintroduce expression of TgPPM3C in the knockout strain (and generate TgPPM3C-COMP strains), sgRNA targeting the multi-stop codon sequence was designed and introduced into a Cas9 plasmid as described above. Donor sequences were designed to restore the original coding sequence of TgPPM3C when co-transfected with the sgRNA used for complementation.

The GRA16 and GRA28 loci were amplified from PruΔku80Δhxgprt genomic DNA with primers to the GRA16 and GRA28 coding sequence and 1.5kb upstream of the start codon, with overhangs to pLIC-3xHA-DHFR sequences. Gibson Assemblies (NEBuilder HiFi DNA Assembly) were subsequently performed to clone GRA16 and GRA28 into PCR amplified pLIC-3xHA-DHFR plasmid backbones. To mutagenize GRA16 serine residues, KLD reactions were performed using the pLIC-GRA16-3xHA plasmid as template DNA for PCR, with primers designed to introduce mutations as intended.

For each transfection, 5×10^6^ to 1×10^7^ Pru*Δku80Δhxgprt* tachyzoites were electroporated in cytomix [50] after harvesting egressed parasites from HFF monolayers and filtering through 5μm filters. Selection of transfected parasites was performed either with 2μM pyrimethamine for at least two passages or with media containing 25μg/mL mycophenolic acid and 50μg/mL xanthine 24 hours post-transfection for 6 days before removing selection media and subcloning by limiting dilution, after sufficient parasite egress was observed. For Cas9 transfections, 7.5μg of uncut Cas9 plasmid and 1.5μg of PCR amplified donor sequence or 280 pmol un-annealed donor sequence were used per transfection. For transient transfection of Pru*Δku80Δhxgprt* parasites with plasmids encoding GRA16-3xHA and GRA28-3xHA, 50μg of uncut plasmid was used. For transfections with mutagenized GRA16-3xHA plasmids, 10μg of uncut plasmid were used for random integration into the genome.

### Immunofluorescence Assays

HFF monolayers were grown to confluency on glass coverslips and infected with egressed tachyzoites at an MOI of 1 for most immunofluorescence assays. For IFN-γ stimulation experiments, infected HFF monolayers were stimulated with 100U/mL recombinant human IFN-γ (R&D Systems) 24 hours post-infection and fixed 6 hours post-stimulation. All coverslips were fixed with 4% PFA for 20 minutes at room temperature, permeabilized in a 0.2% Triton X-100, 0.1% glycine solution for 20 minutes at room temperature, rinsed with PBS, and blocked in 1% BSA for either 1 hour at room temperature or at 4°C overnight. Coverslips were labeled with antibodies as follows: HA-tagged proteins were detected with rat anti-HA 3F10 (Sigma 1:200), parasite cyst wall and parasite cytoplasm by in-house mouse SalmonE anti-CST1 (1:500) and rabbit anti-TgALD1 (1:500, kind gift from Dr. Kentaro Kato) respectively, host c-Myc with rabbit anti-c-Myc (D84C12, Cell Signaling, 1:250), IRF1 by rabbit anti-IRF1 (D5E4, Cell Signaling, 1:500), tachyzoite SAG1 by mouse anti-SAG1 (Thermo Fisher, 1:500). Secondary antibodies conjugated to Alexa Fluorophores 488, 555, 594, and 633 targeting each primary antibody species were used at a dilution of 1:1000 (Thermo Fisher). DAPI counterstain was used to label parasite and host cell nuclei (1:2000). Coverslips were mounted in ProLong Gold Anti-Fade Reagent (Thermo) and imaged using either a Leica SP8 confocal microscope, a Nikon Eclipse widefield fluorescent microscope (Diaphot-300), or a Pannoramic 250 Flash III Automated Slide Scanner (3D Histech).

### Quantitative Image Analysis

For all fluorescence quantification, at least 20 randomly selected fields of view were exported from CaseViewer (3D Histech) software after acquiring images with a Pannoramic 250 Automated Slide Scanner. Identical exposure times were used to detect fluorescence intensity from a signal of interest from each replicate. Exported images were analyzed in ImageJ (NIH), segmenting regions of interest (ROI) manually from fibroblasts that contained individual parasitophorous vacuoles. The mean gray value was measured from each ROI in the fluorescent channel used to detect the signal of interest. Mean gray values were plotted using PRISM 8 (GraphPad). As each mean gray value dataset did not exhibit a normal distribution, nonparametric Kruskal-Wallis tests and Dunn’s multiple comparisons test were used to compare the means between groups to determine statistically significance with PRISM 8 software (GraphPad).

### Immunoblotting

Protein lysates were prepared in radioimmunopreciptation assay (RIPA) buffer after harvesting protein from infected fibroblasts cultures 24 hours post-infection. Laemmli sample buffer was added to samples and boiled for 5min before loading on SDS-PAGE 4-20% pre-cast gradient gels (TGX). Transfer to PVDF membranes (Millipore) was performed in Towbin buffer (20% methanol, Tris/Glycine) for 2 hours at 100V, and blocking in 5% BSA/TBST was performed overnight in 4°C. Membranes were labeled in 5% BSA/TBST with either anti-HA peroxidase conjugated antibodies (Sigma, 1:200) or rabbit TgALD1 antibody (1:200) and anti-rabbit HRP antibodies (Thermo Fisher, 1:10000) followed by development of signal with West Pico Plus Chemiluminescent substrate. Images of labeled blots were collected with a Li-COR instrument (Odyssey Imaging System).

### Plaque Assays

Parasites were harvested from host cells with a 27G needle and filtered through a 5μm filter to remove host cell debris. Parasite numbers were counted with a hemocytometer, and 100 parasites from each strain were added in triplicate to wells containing confluent HFFs in 6-well dishes. Parasites were grown for 14 days before fixing and staining with a 20% methanol-0.5% crystal violet solution. Plaque size was quantified using ImageJ, and Kruskal-Wallis tests with Dunn’s multiple comparisons were performed to test for significance with PRISM 8. Plaque assays were repeated three times, using different batches of parasites and host cells for each experiment.

### Electron Microscopy

For transmission electron microscopy, samples were prepared from human fibroblast monolayers infected with parasites grown under tachyzoite growth conditions for 32 hours. Cultures were fixed with 2.5% glutaraldehyde, 2% paraformaldehyde in 0.1 M sodium cacodylate buffer, post-fixed with 1% osmium tetroxide followed by 2% uranyl acetate, dehydrated through a graded series of ethanol and embedded in L×112 resin (LADD Research Industries, Burlington VT). Ultrathin sections were cut on a Leica Ultracut UC7, stained with uranyl acetate followed by lead citrate and viewed on a JEOL 1400EX transmission electron microscope at 80kv.

### Co-Immunoprecipitation

In two separate experiments, 15cm diameter cell culture dishes containing confluent HFF monolayers were infected at an MOI of 3 with either TgPPM3C-HA or control PruQ non-HA tagged parasites. Dishes were washed with ice cold PBS 24 hours post-infection and lifted off each dish with a cell scraper in 1mL ice cold lysis buffer (50mM Tris pH 7.4, 200mM NaCl, 1% Triton X-100, and 0.5% CHAPS) supplemented with cOmplete EDTA-free protease inhibitor, (Sigma) and phosphatase inhibitors (5mM NaF, 2mM activated Na_3_VO_4_). Scraped cultures were passed through a 27G needle five times and sonicated for 30 seconds total (20% amplitude, 1 second pulses). Sonicated samples were incubated on ice for 30min, supernatant cleared by centrifugation (1000xg, 10min), and incubated overnight in a 4°C rotator with 0.25mg anti-HA magnetic beads (100uL slurry, Thermo Fisher). Following overnight incubation, beads were separated on a magnetic stand and washed twice in lysis buffer and four times in wash buffer (50mM Tris pH 7.4, 300mM NaCl, 0.1% Triton X-100) prior to elution in Laemmli buffer with 50mM DTT, boiling beads for 5min prior to magnetic separation and collection of eluate. Eluates were loaded, washed, and digested into peptides with 1μg of trypsin on S-TRAP micro columns (Protifi) per manufacturer guidelines. S-TRAP peptide eluates were concentrated with a speed vac, desalted in HLB resin (Waters), and concentrated in a speed vac once more prior to running through LC-MS/MS.

### Phosphoproteome Preparation

15cm diameter cell culture dishes containing confluent HFF monolayers were infected in triplicate at an MOI of 5 with either TgPPM3C-HA or ΔTgPPM3C parasites. Dishes were washed with ice cold PBS 36 hours post-infection and lifted off each dish with a cell scraper in ice cold PBS supplemented with phosphatase inhibitors (5mM NaF, 2mM activated Na_3_VO_4_). Scraped cultures were pelleted at 3000rpm for 10min at 4°C. Pellets were resuspended in 500μL S-Trap lysis buffer (5% SDS, 50mM TEAB, pH 7.5) supplemented with cOmplete EDTA-free protease inhibitor (Sigma) and HALT phosphatase inhibitor cocktail (Thermo Fisher). Lysates were sonicated (30 seconds, 20% amplitude, 1 second pulses) and cleared by centrifugation (13,000xg, 4°C). Protein from the supernatant was quantified with a Micro BCA assay kit (Thermo Fisher) and 300μg of protein was loaded, washed, and digested with 3μg of trypsin on S-Trap mini columns (Protifi) per manufacturer guidelines. Phosphopeptides were enriched from 100μg of S-Trap peptide eluates using titanium dioxide beads (TiO_2_, GL Sciences), as previously described [51]. Flow through from titanium dioxide bead enrichment were also collected for LC-MS/MS acquisition and normalization of phosphopeptide abundance by non-phosphopeptides. Following TiO_2_ enrichment, peptides were concentrated with a speed vac, desalted in HLB resin (Waters), and concentrated in a speed vac once more prior to running through LC-MS/MS.

### LC-MS/MS Acquisition and Analysis

Samples were resuspended in 10 μl of water + 0.1% TFA and loaded onto a Dionex RSLC Ultimate 300 (Thermo Scientific, San Jose, CA, USA), coupled online with an Orbitrap Fusion Lumos (Thermo Scientific). Chromatographic separation was performed with a two-column system, consisting of a C18 trap cartridge (300 μm ID, 5 mm length) and a picofrit analytical column (75 μm ID, 25 cm length) packed in-house with reversed-phase Repro-Sil Pur C18-AQ 3 μm resin. Peptides were separated using a 120 min gradient from 2-28% buffer-B (buffer-A: 0.1% formic acid, buffer-B: 80% acetonitrile + 0.1% formic acid) at a flow rate of 300 nl/min. The mass spectrometer was set to acquire spectra in a data-dependent acquisition (DDA) mode. Briefly, the full MS scan was set to 300-1200 *m/z* in the orbitrap with a resolution of 120,000 (at 200 *m/z*) and an AGC target of 5×10e5. MS/MS was performed in the ion trap using the top speed mode (2 secs), an AGC target of 10e4 and an HCD collision energy of 30.

Raw files were searched using Proteome Discoverer software (v2.4, Thermo Scientific) using SEQUEST as search engine. We used both the SwissProt human database (updated January 2020) and the Toxoplasma database (Release 44, ME49 proteome obtained from ToxoDB). The search for total proteome included variable modifications of methionine oxidation and N-terminal acetylation, and fixed modification of carbamidomethyl cysteine. Analysis of the phosphoproteome included carbamidomethylation on cysteine residues as a fixed modification, while phosphorylation on serine, threonine and tyrosine residues was set as variable modification. Trypsin was specified as the digestive enzyme. Mass tolerance was set to 10 pm for precursor ions and 0.2 Da for product ions. Peptide and protein false discovery rate was set to 1%.

Each analysis was performed with either two biological replicates (Co-IP) or three technical replicates (phosphoproteome). Prior statistics, proteins and phosphopeptides were log2 transformed, normalized by the average value of each sample and missing values were imputed using a normal distribution 2 standard deviations lower than the mean. Phosphopeptides were normalized by protein abundance before assessing statistical changes in relative abundance. Statistical regulation was assessed using heteroscedastic T-test (if p-value < 0.05). Data distribution was assumed to be normal but this was not formally tested. Hypergeometric testing for *Toxoplasma* compartment enrichment was performed with a custom R script (github.com/nataliesilmon/toxotools), updated to include recent annotations and Gene IDs from ToxoDB (Release 47, ME49 genome). Generation of sequence logos was performed with Seq2Logo [52], using 31 amino acid sequence windows from either *Toxoplasma* phosphopeptides calculated as significantly enriched in ΔTgPPM3C samples or randomly selected *Toxoplasma* phosphopeptides detected in both wild-type and ΔTgPPM3C samples.

### Mouse Studies

Eight week old female C57Bl/6 mice (The Jackson Laboratory, Bar Harbor, ME) were infected with 16,000 tachyzoites of each strain intraperitoneally. Mortality was observed daily for 30 days. For cyst burden analysis, brains were collected from mice injected intraperitoneally with 2000 parasites 30 days prior. One brain hemisphere per mouse was homogenized with a Wheaton Potter-Elvehjem Tissue Grinder with a 100-150 μm clearance (ThermoFisher) in PBS and an aliquot of the homogenate was viewed under a epifluorescence microscope (Nikon) to count GFP-positive cysts. Kruskal-Wallis tests and Dunn’s multiple comparisons test were performed to test for significance between groups with PRISM 8. A log-rank test was performed in PRISM to test for statistical significance in Kaplan-Meier survival curves.

### Ethics Statement

All mouse experiments were conducted according to guidelines from the United States Public Health Service Policy on Humane Care and Use of Laboratory Animals. Animals were maintained in an AAALAC-approved facility, and all protocols were approved by the Institutional Care Committee of the Albert Einstein College of Medicine, Bronx, NY (Animal Protocol 20180602; Animal Welfare Assurance no. A3312-01).

## ACKNOWLEDGEMENTS

We thank members of the Weiss lab for their comments, suggestions, and insights in the preparation of this manuscript, with special thanks to Rama R. Yakubu and Jessica Weiselberg. We thank Dr. John Boothroyd and members of his lab for useful insights and experimental suggestions. We thank the Albert Einstein Analytical Imaging Facility, specifically Dr. Vera DesMarais and Andrea Briceno for training on various light microscopes and suggestions for ImageJ analysis, as well as Leslie Gunther-Cummins, Xheni Nishku, and Timothy Mendez for electron microscopy sample preparation and training. We thank the Einstein Laboratory for Macromolecular Analysis and Proteomics for their pivotal assistance with all LC-MS/MS preparations and analysis.

This work was supported by P30CA013330, SIG #1S10OD016214-01A1, and SIG #1S10OD019961-01 (Einstein Analytical Imaging Facility), 1F31AI136401 (J.M.), R01AI134753 (L.M.W.), and R21AI123495 (L.M.W.).

## Author contributions

J.M. and L.M.W. conceived and designed the work; J.M., V.T., T.T. all performed experiments involving parasites; J.T.A. and S.S. prepared samples for LC-MS/MS prep and analyzed LC-MS/MS data; J.M. and L.M.W. curated and analyzed all the data; J.M., S.S., and L.M.W. wrote the paper.

## Supplemental Figures, Tables and Datasets

**Figure S1. No gross differences in parasitophorous vacuole morphology are observed between TgPPM3C-HA and ΔTgPPM3C strains.** Representative transmission electron micrographs of vacuoles formed by either TgPPM3C-HA or ΔTgPPM3C parasites, 32 hours post-infection in human fibroblast monolayers. The intravacuolar network, a hallmark of tachyzoite *Toxoplasma* vacuoles, is evident in the vacuoles formed by both strains.

**Figure S2. TgPPM3C protein alignment demonstrates high conservation across *Toxoplasma* lineages.** Clustal Omega alignment of TgPPM3C amino acid sequences from the Type I reference strain GT1, Type II reference strain ME49, and Type III strain VEG. Sequences were obtained from ToxoDB.

**Figure S3. TgPPM3C protein alignment across coccidian relatives demonstrates high conservation in *Neospora caninum* and *Hammondia hammondi***. Clustal Omega alignment of TgPPM3C amino acid sequences from *Toxoplasma* strain ME49 (TGME49_270320), *Hammondia hommondi* strain H.H.34 (HHA_270320), *Neospora caninum* Liverpool (NCLIV_036340), *Sarcocystis neurona* SN3 (SN3_02500075), and *Eimeria acervuline* Houghton (EAH_00048430). Sequences were obtained from EuPathDB.

**Dataset S1. LC-MS/MS data obtained from Co-IP experiments.**

**Dataset S2. LC-MS/MS data obtained from phosphoproteome preps of TgPPM3C-HA and ΔTgPPM3C cultures.**

